# Computational metabolic modeling unveils gut microbiome’s role in metabolic shifts during murine cancer cachexia

**DOI:** 10.1101/2024.09.13.612865

**Authors:** Torben Kuehnast, Isabella Pototschnig, Manuela Träger, Christian Diener, Hansjörg Habisch, Andrea Vogel, Lisa Wink, Alexander Mahnert, Tobias Madl, Thomas Weichhart, Martina Schweiger, Christine Moissl-Eichinger

## Abstract

Cancer cachexia is a multifactorial syndrome characterized by involuntary weight loss, muscle wasting, systemic inflammation, and metabolic alterations, affecting up to 87% of pancreatic and gastric cancer patients. Unlike simple starvation, cachexia is driven by metabolic disruption involving both host physiology and the gut microbiome. While microbiome changes in cachexia have been documented, a coherent understanding of how these changes translate into functional metabolic shifts remains elusive.

In this study, we combined in vivo fecal and plasma metabolomic analyses with a novel computational microbiome simulation pipeline to identify cachexia-associated microbial metabolites. Using the murine MCA207 tumor line and its cachectic derivative CHX207, we differentiated microbiome changes driven by cachexia from those induced by tumor growth. Our computational tool, McMurGut, a murine-tailored extension of MICOM, enabled simulation of microbial metabolic interactions specific to the mouse microbiome, covering 91% of identified genera.

We identified significant abundance changes in 35 microbial genera and corresponding shifts in metabolite production, including reductions in short-chain fatty acids (SCFAs) like acetate and butyrate, alongside increased production of galactose, formate, and propionate. Notably, decreases in SCFA production, particularly by genera such as *Faecalibaculum* and *Dubosiella*, correlated with exacerbated cachectic symptoms. Additionally, the elevated production of formate and galactose, primarily by *Bacteroides* and *Lactobacillus*, suggested altered fermentation pathways in cachexia, potentially linked to increased mucus degradation.

Validation of our computational predictions via NMR metabolomics highlighted key congruencies between predicted and experimentally observed metabolites, supporting the role of microbiome-driven metabolic shifts in cachexia pathology. These findings provide crucial insights into the microbiome’s involvement in cachexia and suggest future avenues for therapeutic interventions aimed at modulating microbial taxa and their metabolic outputs to improve patient outcomes.

## Introduction

Cancer cachexia is a complex and multifactorial syndrome characterized by involuntary weight loss, muscle wasting, systemic inflammation, and metabolic alterations, commonly seen in advanced cancer patients ^1^. Up to 87% of individuals with pancreatic and gastric cancer may experience cachexia ^2^. One in five cancer-related deaths can be attributed to cachexia. Unlike simple starvation, cachexia is not fully reversible by nutritional supplementation, as it is driven by deeper, substantial metabolic disruption involving both host physiology and, increasingly recognized, the gut microbiome ^3,4^.

Cachexia is associated with a disrupted gut barrier integrity along with increased gut permeability and systemic inflammation^3^. While mucus thickness seemed to remain stable in cachectic mice, upregulation of Muc2 (mucus component) and ZO-1 (epithelial tight junctions), along with elevated plasma levels of lipopolysaccharide-binding protein (LBP) and IL-6, reflect substantial gut dysfunction and inflammatory responses^5^. This inflammation, mediated by cytokines such as IL-6, IL-1, and TNF-alpha, accelerates muscle wasting, with recent evidence showing that IL-6 blockade can attenuate cachexia in animal models ^6^.

The gut microbiome plays a crucial role in regulating metabolism and immune responses^7^. In cachexia, specific alterations in the microbiome have been documented previously^8,9^. Among others, *Lactobacillales* was reported to be decreased, while pro-inflammatory taxa like *Enterobacteriaceae* and *Parabacteroides* were increased^1,8,10,11^. In other murine experiments, slightly altered taxonomic changes were observed, describing a decrease of *Romboutsia* and *Lactococcus* and an increase of *Raoultibacter* and *Streptococcus*^12^. However, despite these findings, a coherent understanding of taxonomic changes in cachexia is still lacking. Critically, the focus must shift toward understanding the metabolic interactions mediated by these microbiota, as microbial metabolites offer a more direct link to disease pathology.

At the host metabolic level, cachexia is characterized by a complex upheaval of concentrations and compositions. In the murine plasma, among many others, amino acids, vitamins, and energy-related metabolites such as glucose, citrate and succinate were reported to be significantly reduced^8,13,14^. Glucose and glycogen were significantly reduced in the liver, while muscle reactive oxygen species (ROS) was increasing^13^. Additionally, reductions in short-chain fatty acids (SCFAs) like acetate and butyrate have been linked to the dysbiotic gut microbiota observed in cachexia^12^.

Given the complex and yet incomplete picture of cachexia and the involvement of a multitude of factors, such as altered environmental factors, genera, enzymatic capacities and metabolites, elucidation of functional connections on metabolite and taxonomic levels requires a novel approach that clearly outlines pathways from start to end.

To address this challenge, we combined *in vivo* fecal and blood metabolomic analyses with a novel computational microbiome simulation pipeline to identify metabolites differentially abundant in cachexia, non-cachexia cancer, and healthy controls. Given the absence of a suitable murine microbiome simulation model, we developed McMurGut, a comprehensive metabolic model catalog encompassing 91% of all identified microbial genera. Integrated within MICOM, a mathematical modeling framework capable of simulating microbial growth rates and metabolic activity in the gut, McMurGut enabled us to predict the metabolites produced by the gut microbiota under cachexia and control conditions.

Validation through untargeted NMR metabolomics revealed a remarkable congruence between predicted and experimentally observed metabolites, highlighting key metabolic shifts and the microbial taxa involved in cancer cachexia. These findings provide critical insights into the microbiome’s role in driving the metabolic alterations seen in cachexia.

Future directions may focus on targeted modulation of microbial taxa and dietary interventions aimed at improving metabolic health and enhancing quality of life for cancer patients.

## Results

### Cachexia onset drives distinct microbiome alterations in late-stage tumor progression

The 3-methyl-cholanthrene (MCA)-induced fibrosarcoma cell line MCA207 is a tumor model used to study fibrosarcoma tumor progression in C57BL/6 mice. Due to genetic and phenotypic transition upon passaging and freeze-thaw cycles, the MCA207 cell line evolved to a cachexigenic sub-cell type CHX207^6^. Pototschnig et al 2022 provided a detailed characterization and comparison of MCA207- and CHX207 cancer cells and tumor-bearing mice, focusing on metabolic and immunophenotypic alterations of lean and adipose tissue comparing a non-cachexigenic and a cachexigenic cancer type^6^. In this study, we extended previous work by investigating a cachexia-host-microbiome-interaction. Fecal samples were collected at multiple time points from day 1 to day 13 after cancer cell injection (Fig. 1A) and were analyzed by microbiome sequencing and analysis, *in silico* metabolic prediction, and NMR-based metabolomics of blood and fecal samples. Three independent experiments were conducted, comprising cachexia-groups (CHX207-bearing mice) and control groups (MCA207-bearing mice and PBS-injected mice).

**Figure 1.**
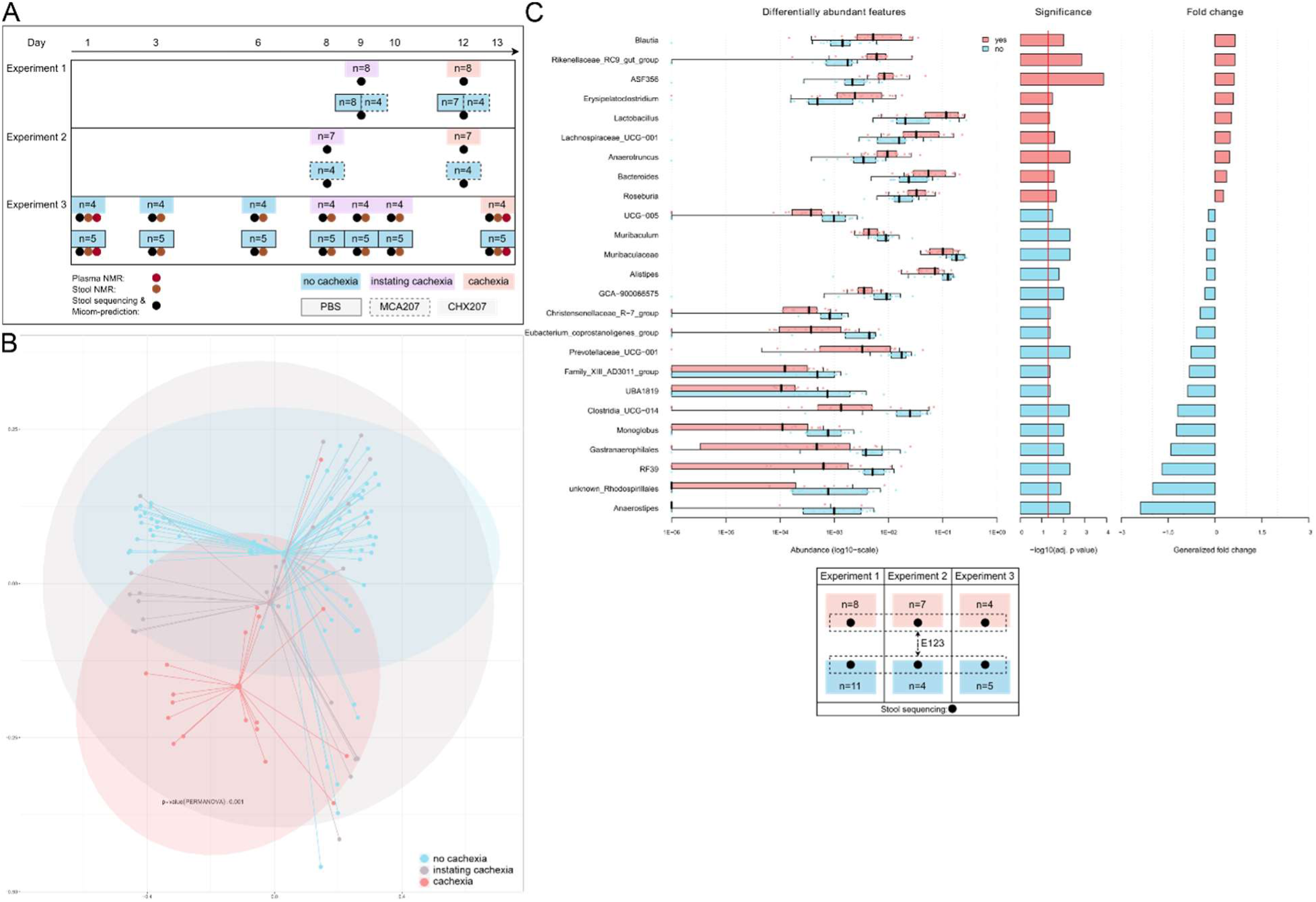
Effects of cachexia on microbial composition. A) Overview of mouse experiments, sampling times and sampling methods, B) Effect of cachexia on microbial composition. Cachexia samples (day 12/13, cachexia group, red) were compared to control samples (controls and CHX-samples until day 6, blue) and instating cachexia samples (purple) (n=124). C) Differentially abundant genera in cachexia. A Wilcoxon test was performed for E123, comparing cachexia samples (day 12/13) to control group (day 12/13). Graph shows top 25 genera (p < 0.05) significantly changed in cachexia. B+C) Panels created in Namco ^15^ and modified in Inkscape.

Cachexia-driven body weight loss could be observed in CHX207-bearing mice, but not MCA207-bearing mice, after day 9, and became statistically significant by day 13 (ANOVA followed by Tukey’s post hoc analysis, p≤0.01). Based on these observations, we categorized the samples into three phases: Early (low-key) cachexia: days 1-6, instating cachexia: days 8-12, and late-stage cachexia: day 12/13.

To determine whether the non-cachexigenic MCA207 tumor affected the gastrointestinal microbiome we analyzed beta diversity using Bray-Curtis dissimilarity. PCoA analysis showed overlapping clusters for the MCA207 and the PBS control group, indicating no significant differences (p = 0.158). Given the similar microbiome parameters (non-significant differences in alpha or beta diversity), we concluded that MCA207 tumors had no substantial effects on the host’s microbiome composition. Consequently, we combined MCA207 and PBS-group as controls for subsequent analyses to focus on the cachexia-specific impact on microbiome composition in CHX207-bearing mice.

To elucidate the effect of the cachexigenic CHX207 tumor on gut microbiome composition, beta diversity was analyzed and compared between samples unaffected by cachexia (controls and CHX-samples until day 6), instating cachexia and late-stage cachexia. Late-stage cachexia samples (d13) showed a significant (p = 0.001, PERMANOVA) change in microbiome composition compared to control samples (Fig. 1B).

Differential abundance analysis (non-parametric Wilcoxon test) revealed significant shifts in the abundance of specific microbial taxa during cachexia (d12/13) (Figure 1C), featuring 10 genera with p<0.05 and 25 with FDR-corrected q<0.05. Genera *Blautia*, *Rikenellaceae*_RC9_gut_group, ASF356 (Lachnospiraceae family), *Erysipelatoclostridium* and *Lactobacillus* were significantly increased, while *Anaerostipes*, Rhodospirillales, RF39 (Bacilli), *Gastranaerophilales*, *Monoglobus* and Clostridia_UCG-014 (Clostridia) were significantly reduced. Among these, *Blautia* showed the highest fold-change and statistical significance (FDR adjusted p-value = 0.0096), marking it as the most prominent taxon favorably influenced by the cachectic gut environment. *Blautia* species are strictly anaerobic and utilize CO, H_2_/CO_2_ and carbohydrates as energy sources, producing acetic acid, succinic acid, and various other metabolites^16^. In contrast, *Anaerostipes* was significantly affected by cachexia (FDR adjusted p-value = 0.0049). Known for their ability to produce butyrate from carbohydrates, *Anaerostipes* representatives have immunosuppressive and anti-inflammatory functions^17^.

### Cachexia drives distinct redistribution of metabolites between gut and plasma during late-stage progression

Samples from experiment 3 (E3) were collected on days 1, 3, 6, 8, 9, 10 and 13 for both 16S rRNA gene amplicon sequencing and fecal NMR spectroscopy (Figure 1A). To minimize health risks for the mice, plasma samples were taken only on days 1 and 13.

Metabolomic analyses revealed that several metabolites - including formate, citrate, tryptophan, uracil, 2-hydroxybutyrate, lactic acid, histidine, glycerol, phenylalanine, alanine and indoxyl sulfate - were elevated in cachexia plasma but reduced or unchanged in feces (Fig. 2). This suggests a cachexia-driven enrichment in the blood and depletion from the gut lumen. Interestingly, metabolites reduced in the plasma (hypoxanthine, acetate, valine, isoleucine, leucine, glucose, lysine, glutamine) were also consistently reduced in the fecal samples. These metabolites may have been transported from the gut to the body, where they were rapidly metabolized or absorbed by tissues and/ or the tumor, rather than accumulating in the blood. Generally, these results indicate a broad, cachexia-driven transfer of metabolites from the gut to the host for immediate use. Notably, no metabolites elevated in the feces showed corresponding changes in the plasma. Metabolites unaffected by this host-driven pull, and universally increased in feces, included galactose, isopropyl alcohol, trimethylamine, asparagine and 5-aminopentanoic acid.

**Figure 2.**
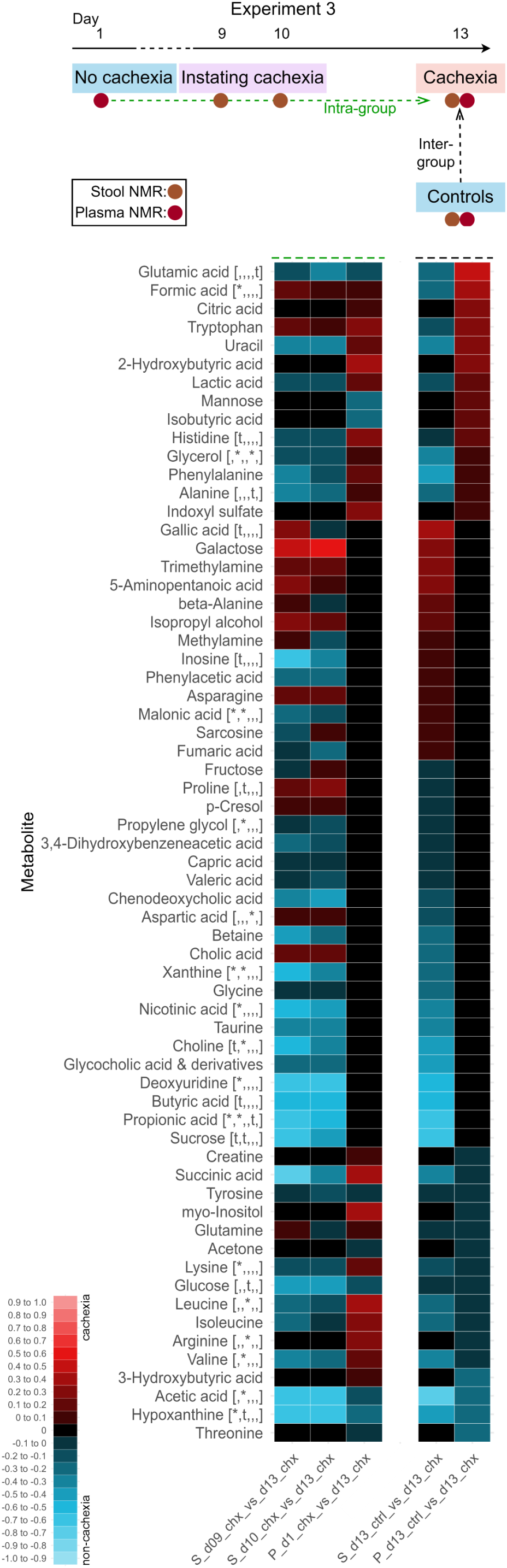
**Differentially abundant metabolites of the stool (S) and plasma (P) in cachexia**, shown as fold-changes of the means. Comparing instating (day 9 / day 10) to late-stage cachexia (day 13) (columns 1 and 2), and d13 controls to late-stage cachexia (column 4) in stool; and initial (day 1) to late-stage cachexia (day 13) (columns 1) and d13 controls to late-stage cachexia (column 2) in plasma, all in experiment 3. Sorted by column 5. Red indicates increase in cachexia, cyan indicates reduction in cachexia. Statistics in square brackets [column 1, column 2, column 3, column 4, column 5] with * = p < 0.05 and t = trend of p between 0.05 and 0.1, based on paired t-test (column 1, 2, 3, 5) and unpaired Welch t-test (column 4). Data obtained by NMR spectroscopy. Normalization of fold-changes were calculated by multiplying values below 1 with 1/x*-1 and fold-changes above 1 were left unchanged. 1 was subtracted from positive values and added to negative values.

### Development of McMurGut 1.1 for precise metabolic modeling of the murine gut microbiome

To enhance traditional microbiome analyses, and to link microbiome observations with experimental metabolome data, we aimed to simulate the metabolic network based on the given microbiome composition *in silico*. For this, we utilized MICOM (Metagenome-Scale Modeling to Infer Metabolic Interactions), an established mathematical modeling framework ^18,19^. While originally created to simulate metabolic interactions within human gut microbial communities, a functional workflow for the murine gut microbiome was still lacking. This gap is substantial, as many mechanistic studies in the field rely on murine models. To address this, we created McMurGut 1.1, a metabolic model catalog for the murine gut and implemented it into MICOM (Fig. 3). McMurGut is fully customizable and contains: i) a comprehensive catalog of genome-derived metabolic models of individual species, covering up to 91% of all taxa detected in murine fecal samples from this study, and ii) a “growth medium” for the simulations, which reflects the composition of standard murine chow and specific features of the mouse gut environment, such as mucin.

**Figure 3.**
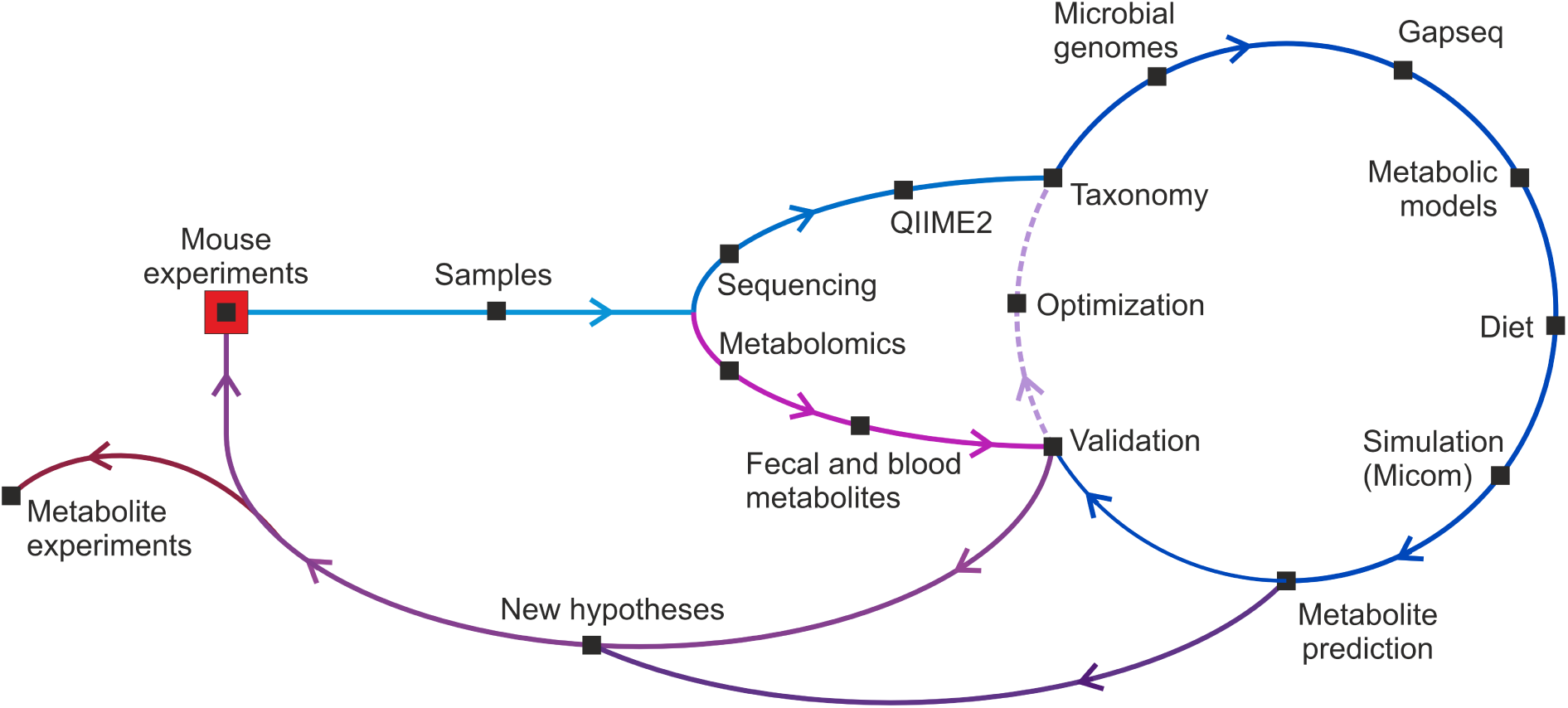
Overview of the McMurGut pipeline. Samples from mouse experiments were processed via sequencing and combined to a list of taxa. After screening for respective genomes, Gapseq was used to create metabolic models. Together with a newly created mouse medium, metabolites were predicted via Micom. Candidates were cross-validated to NMR-spectroscopy, followed by rounds of optimization and repetitions. Finally, validated metabolites were used for generating new hypotheses on functional metabolite pathways in cachexia.

The mathematical framework of choice (MICOM) relies on annotated genomes of taxa, reflecting their individual metabolic capacity at the enzyme, pathway and transporter levels^18^. Initial tests using the available AGORA2 database^20^ and the recently published unified genome catalog of genomes from the murine gut (MGnify)^21^ revealed limited coverage, with only 68% and 51% of the taxa found in our murine samples. To avoid a severe loss in prediction accuracy, we developed a customized approach through McMurGut. We searched for all annotatable genera (conforming to hierarchical nomenclature) in our mouse experiments, identified all species described so far for each genus and collected (if available) three representative genomes for each of the species. This was done by prioritizing murine-specific (MGBC^22^, CMMG^23^) and supplementing missing genomes with those from UHGG^24^ and NCBI. The selected genomes were converted into model reconstructions using gapseq, a tool which is based on a curated reaction database and advanced gap-filling algorithm^25^.

A total of 754 models were created and incorporated in McMurGut 1.1, representing 134 taxonomically classifiable genera. This resulted in 91.1 % coverage of all genera (123 out of 135) detected in experiments E123. In E3, a similar coverage of 91.8 % (123/134) of all genera was achieved. Importantly, the 91.1% coverage in E123 corresponded to 98.4% of all taxonomically annotatable reads (4,252,236 out of 4,321,933 reads).

Although these high coverage rates provide confidence in the accuracy of upcoming simulations, it is important to note that a substantial portion of reads (36.9% of total) could not be taxonomically annotated at genus level and thus could not be included. The retrieval of genomes from these uncovered genera and the placement of these in genome databases may erase this issue over time.

Simulations in MICOM require nutrient input fluxes, reflecting the molecules available to the microbial models. We provided this by creating a diet table (“medium”) that contained all available metabolites to the gut microbiome along with their respective quantity and flux (mmol per hour). While various nutrient compositions for various human diets are readily available, none existed for mice. To create a custom version for our lab mice, we converted the mouse chow (by ssniff Spezialdiäten GmbH, Germany) from percentage or weight into mmol per hour. Additionally, mucus, murine bile acids and water were added to the formula. To ensure completeness, we used MICOM’s medium completion function, which retroactively performed a growth simulation of the McMurGut 1.1 models using the ssniff-based diet draft. If a model failed to grow, essential metabolites were added in minimal quantities to match the predicted enzymatic pathways of the individual models. This approach resulted in a fully complemented diet file (“growth medium”), enabling 131 of 132 genera to achieve a growth rate above.

### Tradeoff optimization and validation of McMurGut 1.1 metabolic predictions against *in vivo* experimental data

Although MICOM has previously demonstrated accurate predictions of metabolic production in human studies^19^, a thorough calibration of the available parameters and inputs has been proven critical for the development of this work. During tests with alpha versions, we identified several factors that negatively impacted prediction accuracy, including the use of an unsuitable diet (e.g. a Western human diet), outdated versions of MICOM and gapseq, and lower genus coverages.

To benchmark the quality and accuracy of the newest McMurGut 1.1 version, we first defined MICOM’s tradeoff between community growth and individual strain growth. We then compared NMR measurements from stool and plasma to the metabolic predictions. As all relevant measurements were available only in experiment E3, we focused on day 13 (cachexia) samples, comparing them to the control samples from the same day. This allowed us to evaluate the accuracy of predicted metabolic changes based on the taxonomic profile derived from microbiome analyses.

In MICOM, a cooperative tradeoff can be chosen which tries to balance community growth and individual strain growth. With a low value, the system is set to prioritize growth of a few taxa, which may be important in research of dominant species versus dormant ones. A high value allows similar growth among all members, keeping the balance (Figure 4A). Generally, a higher value is considered more realistic for an interacting community, as it may reveal synergistic metabolic networks. For this, however, the medium composition must be sufficient to allow it.

**Figure 4.**
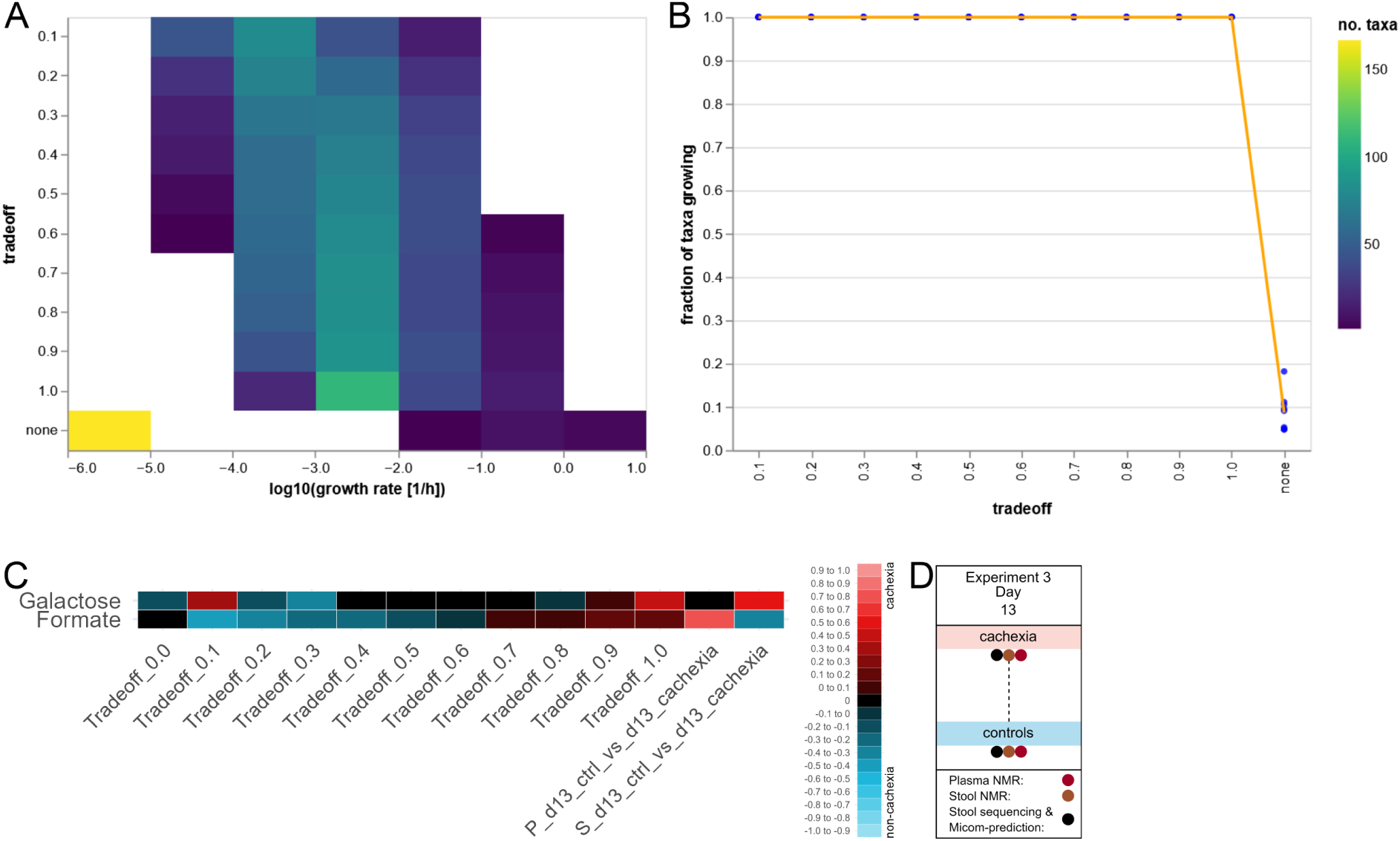
Tradeoff analysis. In silico analyses run via Micom (v0.35.0), MCMG754 and mouse chow diet. Comparisons based on D) E3 at day 13, cachexia vs control. A) Predicted growth under varying tradeoff conditions from 0.0 to 1.0, showing the amount of species growing under respective conditions with specific growth speed. B) Total number of species growing in principle under various tradeoffs. A+B) Created with Micom’s plot_tradeoff function. C) Metabolic prediction, plasma and stool NMR of metabolites increased (red) or decreased in cachexia depending on tradeoff parameter, compared to NMR of galactose and formate in plasma (P) and stool (S). For each method (plasma, stool, prediction) and direction (increase, decrease), the maximum of the respective value-group (of all detected metabolites) was normalized to 1 or -1 (depending on increase or decrease), with the others ranging between 1 and 0, or -1 and 0 respectively. Increased microbial production from both metabolites was best reflected by tradeoff 1.0.

To determine the highest (optimal) tradeoff value for experiment E3 (day13 cachexia vs day13 controls) at which all taxa were able to grow, MICOM’s tradeoff() function was used. Across all tradeoff values tested (from 0.1 to 1.0), all taxa were able to grow (Figure 4B), as the curve remained at 1.0 (100%). This result was expected, as we specifically tailored the diet file to meet the demands of the individual McMurGut models. Based on these findings, a tradeoff value of 1.0 was considered the most realistic for simulating the metabolic network.

It is important to note that for the validation of the simulation against experimentally derived in vivo data, all methods used - 16S rRNA gene sequencing for microbial profiling and NMR metabolomics (Figure 1A) - were performed on the same fecal samples (with one half allocated for NMR and the other for DNA extraction). Plasma and fecal samples were collected simultaneously on day 13.

From the list of differentially abundant metabolites, two metabolites - formate and galactose - were of particular interest and used for trade-off validation. Both showed a marked increase in either plasma or stool samples (Fig. 2), and are recognized products of the gut microbiome. Formate is a commonly known, highly abundant product of microbial fermentation. Although formate can also be produced by the host through the degradation of serine^26,27^, serine concentrations remained stable throughout the experiment, reinforcing the likelihood that the elevated formate originated from microbial activity. Galactose, found to be differentially abundant in the gut during cachexia, is produced by microbial activity, typically through the cleavage of lactose^28^. With a tradeoff factor of 1.0 (i.e. the maximum growth of all microorganisms) we observed the highest production of formate and galactose by the microbiome, aligning with the experimental NMR data, showing an increase of galactose in the stool samples and an increase of formate in blood under cachexia conditions (Fig. 4C). Based on these results, a tradeoff factor of 1.0 was chosen for all further calculations.

### *In silico* modeling identifies critical metabolites and key microbial contributors

In the next step, we applied the metabolic prediction method, integrating McMurGut and MICOM, to the combined dataset from all three independent cachexia experiments (Fig. 5). We predicted several metabolite candidates to be associated with the cachectic microbiome. Notably galactose, GABA, spermidine, formate, propionate and others were elevated, while pantothenic acid, butyrate, acetate, succinate and fumarate were decreased.

**Figure 5.**
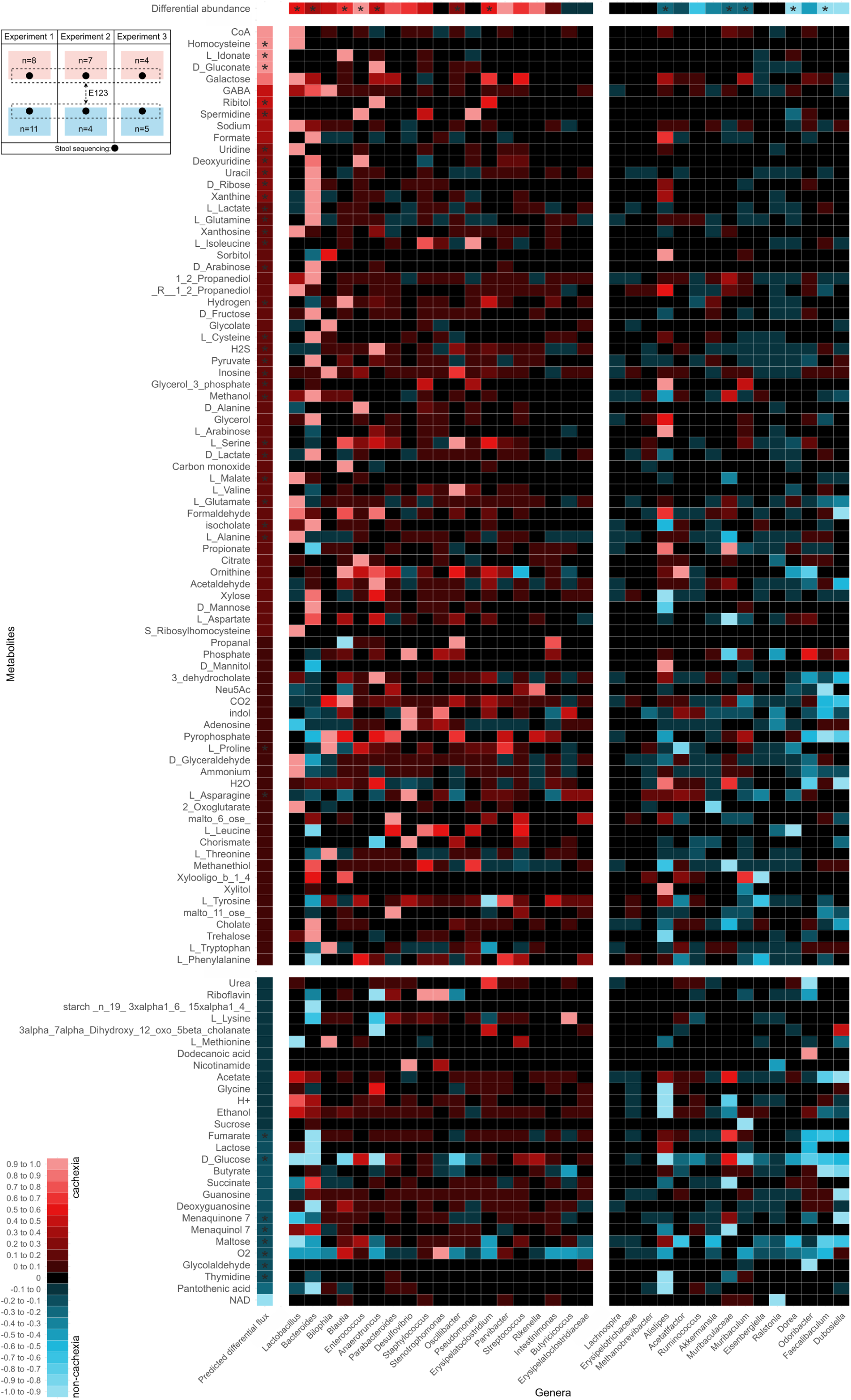
Predicted differential metabolite production in cachexia. Heatmap shows differential abundance of metabolites and genera, comparing late-stage cachectic samples to control samples (E123). Colors change in 0.1 relative steps to maximum of 1, with red = increased in cachexia, blue = increased in controls and black = zero or no value. Deprotonated and protonated metabolites were treated tautologically. Sorted by the mean prediction of each metabolite on the y-axis (predicted differential flux), and the sum of metabolite production of each genus on the x-axis (not shown), leading to an accumulation of cachexia-associated metabolite-genus combinations in the top-left. **Predicted differential flux**. Prediction based on Micom 0.35.0, tradeoff=1.0, McMurGut 1.1 (MCMG754 + completed mouse chow diet), E123, at day 12/13 cachexia vs d12/13 control. The results of compare_group [log2(mean(sum(flux*abundances per metabolite and cachectic sample))) minus log2(mean(sum(flux*abundances per metabolite and control sample)))] were normalized to the 95th quantile of all (absolute) values of the same metabolite to be in the range of -1 to 1. Statistics for metabolites was added with * for p < 0.05, based on Mann-Whitney U test. **Differential abundance of genera in cachexia**. Based on differential abundance associations calculated with Namco^15^, analyzing E123 at day 12/13 cachexia vs day 12/13 controls. Significantly different abundances were calculated with non-parametric Wilcoxon test at days 12/13 (n=39). * = p-value < 0.05. The fold-changes of genera were not normalized. **Individual metabolite production per genus in cachexia.** Difference of the sum of flux*abundance of metabolites produced by individual genera that were differentially produced by metabolites in cachexia vs control. Based on Micom 0.35.0, tradeoff=1.0, McMurGut 1.1 (MCMG754 + completed ssniff diet), containing E3 day 13 cachexia vs d13 control. Fluxes and abundances were extracted after applying Micom’s grow- and build-functions (res[1] dataframe) via search_flux_species.ipynb. The sums of flux*abundances of metabolites per genera and condition were normalized to the (absolute) maximum of all values to be 1, with other values ranging between +-1 and 0.

The advantage of a simulation is that it allows to track metabolic production down to the producing taxa. This was achieved by extracting flux*abundances *from each genus* throughout all samples of the same conditions by multiplying taxon abundance (number of sequence reads) and flux (general capability to produce a metabolite under the given synergistic interactions in the same sample) (Fig. 5).

Lastly, the predicted metabolites for each condition and flux*abundances for each genus and condition were augmented with differential abundances of each genus (Fig. 5), creating a complex map. Following galactose, its mean increase may be attributed to flux*abundance’s concurrent increase in 14 genera, such as *Lactobacillus*, *Erysipelatoclostridium*, *Streptococcus*, *Bacteroides* and *Muribaculum*. Of note, GABA production was observed, possibly associated with a higher activity/abundance of *Lactobacillus* and *Bacteroides*. Conversely, the reduced butyrate availability was attributed to reduced contributions from *Faecalibaculum*, *Dubosiella*, *Butyricicoccus*, *Oscillibacter* and *Acetatifactor*.

### Experimental observations validate *in silico* predictions of metabolite changes and highlight potential pathways and adaptive responses of the microbiome during cachexia

In order to identify the most crucial candidates (metabolites and taxa) involved in cachexia during cancer progression, which could be most useful for further hypothesis building and subsequent experiments, we concentrated our analysis to those metabolites that overlapped in all metabolic predictions (E123, E3) and were detected in plasma or feces via NMR metabolomics, and their association with specific microbial taxa.

In strong agreement with NMR experimental data, the *in silico* prediction was able to picture the (significant) decrease of succinate, butyrate, acetate, fumarate and glucose (Fig. 6A). This reduction was attributed to a reduced abundance/activity of various microbial genera, such as *Faecalibaculum* and Muribaculaceae, *Dorea*, *Acetifactor*, *Butyricicoccus,* and *Blautia*. All these metabolites are highly critical for the maintenance of host health and gut barrier, and a reduction of these might explain the loss of gut barrier in cachexia patients. In particular butyrate is known to orchestrate macrophage polarization, mitigating muscle atrophy and averting cachexia-induced muscle deterioration^29^.

**Figure 6.**
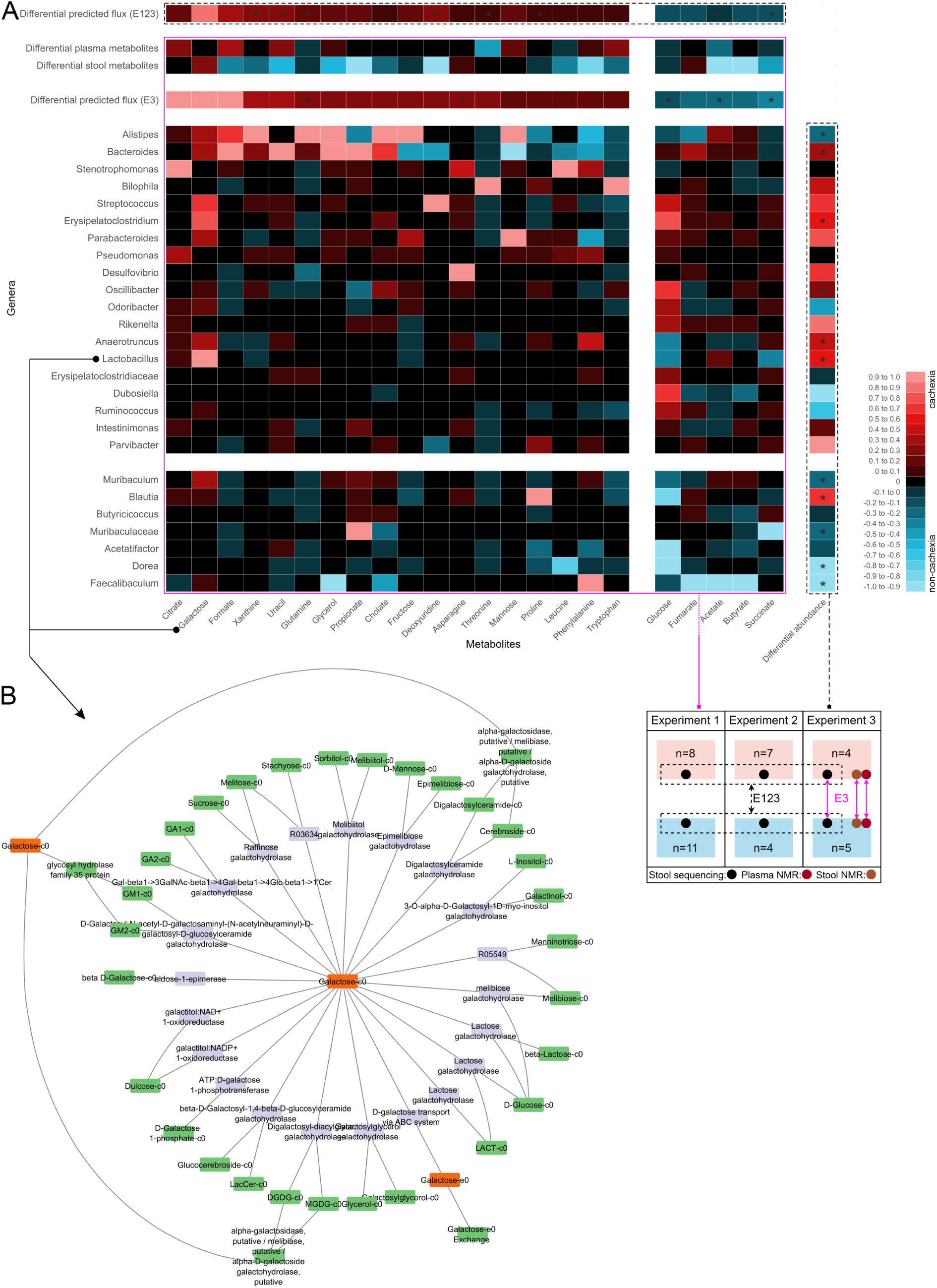
Map of metabolic flux in cachexia. **A)** Heatmap shows differential abundance of metabolites or genera, comparing late-stage cachectic samples to control samples. Colors change in 0.1 relative steps to maximum of 1, with red = increased in cachexia, blue = increased in controls and black = zero or no value. Deprotonated and protonated metabolites coming from prediction or spectroscopy were treated tautologically. Shown are only those metabolites that matched in prediction in E123 (d12d13) and E3 (d13) and appeared in NMR (E3, d13). Sorted by total sum of production of each genus on y-axis, and mean flux*production from all genera of each metabolite (Differential flux predicted (E3)) on x-axis, leading to an accumulation of cachexia-associated metabolite-genus combinations in the top-left. Predictions based on Micom 0.35.0, tradeoff=1.0, McMurGut 1.1 (MCMG754 + completed ssniff diet). Data within the pink square was based on E3 and within the black dotted square was based on E123. (CENTER) Sum of flux*abundance of metabolites produced by individual genera differentially produced in cachexia and from control. The sums were normalized to the (absolute) maximum of all values to be 1, with other values ranging between +-1 and 0. (RIGHT) Fold-changes of genera’s abundance in cachexia or control. Based on differential abundance associations calculated with Namco (Figure 1). Significantly different abundances were calculated with non-parametric Wilcoxon test (n=39). * = p-value < 0.05. (TOP) Change of metabolites in cachexia, either predicted via Micom or detected as fold-change via NMR (stool or plasma). Fold-changes of the predicted mean-fluxes were normalized to the 95th quantile of all (absolute) values of the same metabolite to be in the range of -1 to 1. Fold-changes of NMR-spectroscopy were calculated by multiplying values below 1 with 1/x*-1 and fold-changes above 1 were left unchanged. 1 was subtracted from positive values and added to negative values. Significances for metabolites were indicated by * = p < 0.05, based on Mann-Whitney U test (E123 and E3 prediction), unpaired Welch t-test (stool NMR) and paired t-test (plasma NMR). **B**) Metabolic network of galactose, taken from the gapseq-created model of *Lactobacillus* (genome MGBC000013), visualized via Cytoscape 3.10.2 and focused on galactose’s next two connected nodes. Progenitor metabolites (green) may be involved in galactose (orange) production via specific enzymes (blue).

*Alistipes*, *Bacteroides*, *Stenotrophomonas*, *Bilophila* and *Streptococcus,* showed increased predicted metabolite production during cachexia, in particular by increasing flux*abundance of citrate, galactose, formate, xanthine, uracil, glutamine, glycerol and propionate. This increase aligns with the mean MICOM predictions from the combined dataset E123 and the sum of individual flux*abundance production. NMR spectroscopy confirmed that most of these predicted metabolites indeed increased in either feces *or* the plasma. Galactose, in particular, showed a marked increase across all levels, both in predictions and in feces NMR, indicating a collective rise in its production by the microbiome during cachexia. However, this increase also corresponds with an apparent loss of galactose due to excretion, suggesting that the metabolite is not fully utilized by either the microbiome or the host. This pattern highlights the impact of cachexia on microbiome metabolism, potentially accelerating energy acquisition at the expense of short-chain fatty acid production. It may also involve reduced lactose cleavage in the small intestine and altered liver function.

In contrast to all other metabolites, the genera were consistently predicted to unexceptionally increase their galactose production (Fig. 6A).

Regarding the galactose pathway, *Lactobacillus* appears to be the primary contributor (22.5%), followed closely by *Erysipelatoclostridium* and *Streptococcus*. The respective genome-based models were extracted from McMurGut, comprising seven models for the genus *Lactobacillus* from two species (*Lactobacillus intestinalis* and *Lactobacillus johnsonii*). All seven genomes originated from the Mouse Gastrointestinal Bacterial Catalogue (MGBC)^22^, implying their association to the murine gut. *Lactobacillus intestinalis* (MGBC000013) is displayed as representative for all models (Fig. 6B), showing the same or similar (*Streptococcus*) pathways of galactose production. Therein, galactose may be produced through various hydrolases, which may hint that in the context of cachexia, the microbiome increased its carbohydrate catabolism towards fiber and mucus.

## Discussion

Cachexia, a severe condition marked by substantial weight and muscle loss, significantly impacts cancer patients’ health and survival. As research increasingly focuses on microbiome modulation as a potential therapeutic strategy, it is crucial to elucidate the role of microbiome metabolites in cachexia. Despite numerous studies, results on microbiome metabolite production in cachexia have been inconclusive, often due to technical limitations. To address these gaps, we integrated *in vivo* fecal and blood metabolomic analyses with a novel computational microbiome simulation pipeline for murine models to identify cachexia-related metabolites and their microbial sources.

Our study utilized the MCA207 tumor line and its cachectic derivative, CHX207, which allowed us to differentiate microbiome changes driven by cachexia from those induced by tumor growth. As such, and also by controlling for the cage effect, we were able to set-up a clean experimental procedure, as a basis for our study. We identified 35 microbial genera with a significant abundance change during cachexia (10 genera with p < 0.05, 25 with FDR-corrected q < 0.05, Figure 1C). Specifically, genera from Lachnospiraceae (UCG-004, ASF356), *Enterococcus*, *Blautia*, Rikenellaceae (RC9_gut_group), *Tuzzerella*, *Lactobacillus* and *Anaerotruncus* increased, while *Anaerostipes*, *Monoglobus*, *Dorea*, *Faecalibaculum*, *Tyzzerella*, *Alistepes* and *Muribaculum* decreased in abundance. Overall, these findings align with some previous reports such as those by Jeong et al., and Liu et al. (XXX), though notable discrepancies exist with *Lactobacillus*, which was reported to decrease in some studies but increased in ours (FDR-corrected p-value = 0.044). In detail, Jeong et al reported *Streptococcus*, *Marvinbryantia*, *Candidatus_Arthromitus* and *Erysipelotrichaceae* increasing in cachexia and *Colidextribacter*, *Ruminococcus* and *Romboutsia* decreasing^12^. These genera were in line with ours (exception Clostridium_sensu_stricto_1), but our p-values were above 0.05, indicating the existence of a core group of genera associated with murine cachexia across various studies. Liu et al 2023 showed significant changes in murine cachexia of *Lactobacillus* (increase), *Bacteroides* (Increase), *Enterococcus* (increase) and *Muribaculum* (decrease), again in line with our results. Another study saw differential changes in murine cachexia of *Parabacteroides* (increase, ns^9^) and *Ruminococcaceae* (decrease, ns^10^). However, *Lactobacillus* was reported to decrease and described as potential probiotic in cachexia ^9^.

Our NMR-spectroscopy results showed a marked depletion of various metabolites in feces from cachectic mice, including SCFAs (acetate, butyrate, propionate), deoxyuridine, hypoxanthine and xanthine, sucrose and glucose, nicotinic acid, taurine, succinate, choline and CDCA (Fig. 2). Conversely, some metabolites like galactose, isopropyl alcohol or trimethylamine were elevated. In parallel, several of the metabolites reduced in the fecal samples were elevated in the plasma, such as formate, glutamate, tryptophan, uracil, lactate, histidine and glycerol. This may suggest that during cachexia, the host increased its demand for many of the metabolites, increasing uptake from the gut lumen into the bloodstream.

To better understand these shifts, and with the goal to generate functional hypotheses for the origin and contribution to cachexia, we developed McMurGut, a tailored model for simulating murine microbiome metabolism. As the framework of MICOM was built for the human gut microbiome with human-based selections of microbial models and human-based nutrient data sheets, this framework had to be replaced to fit to the conditions of mouse experiments. These modifications were collected in McMurGut, containing i) a model catalog called MCMG754, including 754 models of strains reflecting 134 genera identified in our mouse experiments, ii) a diet file based on mouse chow, complemented to allow minimum growth for all MCMG754 models. McMurGut featured the ability to individually tailor the model catalog to the targeted microbiome. By this, we could increase the coverage of all genera identified in the cachexia mouse experiments to ∼ 91% (for both, E123 and E3 alone) corresponding to 98.4% of all annotatable reads from sequencing. Best alternative coverage with pre-selected model- or genome catalogs, at this time, was the human model catalog AGORA2, offering a coverage of 68% of all genera^20^. Additionally, we provided to our knowledge the first publicly available lab-mouse diet for microbial metabolite simulation. McMurGut’s validation against NMR data underscores its robustness in simulating microbial metabolite production.

*In vivo* microbiome research often faces challenges in elucidating functional mechanisms behind observed metabolic changes. However, computational simulations offer a powerful advantage, as they allow for detailed tracing of enzymatic reactions and metabolic pathways. By leveraging this capability, we tracked metabolite production back to specific microbial genera, providing insights into the functional roles of these microorganisms in cachexia (Figures 5 and 6). This approach enabled us to identify the final metabolic pathways leading to the production of key metabolites, visualized through metabolic reconstruction models (Figure 6B).

Our simulations revealed that genera such as *Lactobacillus, Bacteroides, Bilophila, Blautia* and *Enterococcus* increased their overall metabolite production during cachexia, while *Dubosiella, Faecalbibaculum, Odoribacter, Dorea* and *Ralstonia* showed reduced production (Figure 5). Interestingly, the altered metabolite production did not necessarily align with shifts in genus abundance. For example, *Stenotrophomonas* increased and *Ralstonia* decreased their total metabolite production, while the abundances of both remained stable. This discrepancy suggests that cachexia induces shifts in microbial metabolic networks, potentially altering or bypassing existing pathways.

Our metabolic predictions indicated a reduced microbial production of vitamins and cofactors (riboflavin [vitamin B2], nicotinamide [B3], pantothenic acid [B5], NAD, menaquinone 7 and menaquinol 7 [vitamin K]). This is in line with previous work in human patients, showing a reduction of vitamins in the plasma of cachexia patients^8^. In theory, symptoms of vitamin deficiency and cancer cachexia overlap, such as fatigue and weakness, impaired metabolic processes, muscle wasting, bone loss, cognitive decline, neurological symptoms, system inflammation and poor wound healing^30,31^. Acetate and butyrate levels were notably reduced in feces, with acetate also decreasing in blood concentration (Fig. 5 and 6). This reduction is in line with previous studies (human: ^11^, mouse: ^10,12,32^) and may be attributed to decreased microbial production. The loss of these SCFAs could contribute to the host’s energy depletion and exacerbation of cachexia.

Our simulations identified key genera involved in shifts in SCFA production. *Muribaculaceae*, *Lactobacillus*, *Bacteroides* and *Alistipes* increased their metabolite production, while *Dubosiella* and *Faecalibaculum* among others, contributed less, leading to an overall reduction. Specifically, the loss of butyrate production was largely due to reduced contributions from *Faecalibaculum*, *Dubosiella* and *Butyricicoccus.* The decline in acetate and butyrate may thus be closely linked to the decreased abundance of these key genera. Acetate is crucial for energy metabolism, as it can be converted to Acetyl-CoA in the liver for various biochemical pathways. Its reduction in cachexia suggests a loss of energy availability for the host. Butyrate has been shown to significantly reduce cachectic effects in mice, such as skeletal muscle atrophy, inflammatory and oxidative stress responses, less macrophage infiltration into skeletal muscles and promotion of M2 macrophage polarization^29^, while serving also as major energy source for colonocytes. Its reduced levels in cachexia could worsen muscle wasting and other cachexia symptoms.

In contrast, other metabolites were found to be increased, such as formate, propionate, and galactose. Formate, produced during fermentation of carbohydrates^33^ or as a side product of choline degradation from TMA to TMAO and formate was elevated in the microbiome, particularly by *Bacteroides*, *Alistipes* and *Stenotrophomonas* (Figure 6A). Despite reduced fecal levels, increased blood concentrations suggest heightened production and absorption, potentially linked to cachexia-related processes. This may be detrimental to the host, as formate was previously associated with colorectal cancer progression^34^. Propionate levels were predicted to be increasingly produced by the microbiome, especially by *Muribaculaceae* and *Bacteroides*, although this compound was found reduced in stool and plasma. This may reflect rapid uptake and utilization by the host. Similarly, increased galactose in feces, without a corresponding rise in plasma levels (Fig. 6A), suggests higher microbiome production with limited absorption into the bloodstream.

Galactose production was notably attributed to *Lactobacillus* (22.5% E3, 23.1% E123) and *Streptococcus* (15.1% E3, 13.8% E123) (Fig. 5 and 6A), which utilize both epithelial epithelial membrane-associated glycolipids and sphingolipids, and non-digestible carbohydrates and oligosaccharides (i.e. raffinose family oligosaccharides (RFOs)) (Figure 6B). It may be hypothesized that in cachexia, the microbiome switches to increased mucus, epithelia surface and fiber degradation. Interestingly, it was previously shown that the mucus-layer in cachexia tended to shrink while mucus production was significantly increased (muc2 and ZO-1)^5^. With this at hand, it may be postulated that in cachexia, the microbiome shifts to an intensified mucus and (if available) fiber fermentation as energy-source, with respective species with these skills increasing in abundance.

In conclusion, our findings provide a comprehensive view of microbiome-related metabolic alterations in cachexia. The integration of metabolomic and computational approaches offers valuable insights into the dynamic interactions between the microbiome and host metabolism in cachexia. Future research should explore these metabolic shifts further to develop targeted interventions that could improve patient outcomes.

## Acknowledgements

Research was supported by funding by the Austrian Science Fund (FWF) Sonderforschungsbereich SFB F83. We are grateful to Daniela Fulterer, for excellent experimental support.

## Methods

### Animal studies

Mice were bred and maintained under specific pathogen-free conditions at housing temperatures of 21-23°C in a 14 h light/10 h dark cycle and fed a standard chow diet with ad libitum access to food and water. C57BL/6J mice were obtained from Janvier Labs, France. Basic details on animal studies including experiment setup and cancer cell culture were previously published by Pototschnig et al^6^. Analysis is based on three independent experiments (E1,2,3). Briefly, 11-week-old (E1) or 12-week-old (E2,3) male C57Bl/6J mice were injected with 1 x 106 non-cachexigenic MCA207 (E1,2) or cachexigenic CHX207 (E1,2,3) fibrosarcoma cells in 100 µl 1 x PBS into the musculus gastrocnemius (m.g.+s.) of the right hind leg. The control group was injected with 100 µl 1 x PBS (E1,3). Feces and blood samples were taken at given time points (see individual experiment setups E1,2,3 for exact schedule). Feces samples were collected directly after defecation, snap-frozen in liquid nitrogen and stored at -80°C until further analysis. For plasma NMR analysis, blood was drawn at the retro-orbital plexus and plasma was obtained from total blood by centrifugation at 1 000 x g and 4 °C for 15 min. In order to gain consistent analysis and insight upon NMR and DNA sequencing, feces samples were thawed, halved on ice-cooling and refrozen at -80°C.

### Metabolomics

Murine feces droppings and blood samples were analyzed via NMR-spectroscopy as previously published^35^.

### Sequencing of 16S rRNA genes

Microbial cells from the halved feces droppings were opened using MagNA Lyser including 100 µl of 0.5 µm beads, alternating 30 seconds of 3500 rpms with 30 seconds on ice for two times total. Supernatant was taken for DNA extraction via QIAamp® PowerFecal® Pro DNA Kit (Qiagen) according to the manual. DNA was eluted in 50 µl of water. Quantity was measured via NanoDrop Spectrophotometer (Thermo Scientific). Aliquots were diluted to total DNA concentration of approximately 10 - 19 ng / µl. 1 µl of this was used as template for PCR, targeting the 16S rRNA gene in the V4-region via primers Illu-515FB_2018 with sequence TCGTCGGCAGCGTCAGATGTGTATAAGAGACAGGTGYCAGCMGCCGCGGTAA and Illu-806RB_2018 with sequence GTCTCGTGGGCTCGGAGATGTGTATAAGAGACAGGGACTACNVGGGTWTCTAAT^36^.

Library preparation and sequencing of the amplicons were carried out at the Core Facility Molecular Biology of the Center for Medical Research at the Medical University Graz, Austria. Sequencing was performed in paired-end run mode on an Illumina MiSeq with v3 600 chemistry and 300 bp read length57. Raw reads were processed using Qiime2 v2022.8. Briefly, reads were imported as paired end sequences, adapters (forward and reverse primers, see above) were trimmed via cutadapt and sequences were denoised with DADA2, truncating at 165 forward and 150 reverse reads. Finally, taxonomy was assigned using pre-trained Naive Bayes classifier based on the SILVA 138 database. Raw reads are available at European Nucleotide Archive PRJEB37883.

### McMurGut 1.1 - MCMG754 creation

Taxonomy files were merged into a list of genera. Entries not clearly associated with a taxonomical genus were discarded. Genome catalogs MGBC, CMMG, UHGG and NCBI were searched for the genera, and (if available) 3 reference genomes for each species present in each genus were downloaded. The collected genomes were compressed to .fna.gz format. CMMG genomes were transformed from .gff.gz into .fna.gz via GFFintoFASTAv06.ipynb. Genomes were used to create metabolic models via gapseq 1.2. ^25^, via gapseq doall. Resulting .XML-files were combined to a catalog via Qiime2’s export tool.

### McMurGut 1.1 - MCMG754 coverage

Taxonomic tables from all mouse experiments were reduced to contain only taxa and abundances from the two compared conditions (i.e. E123 day 12/13 samples or E3 day 12/13). Duplicates and entries that didn’t contain acknowledged genera were removed. The resulting lists were compared to the list of genera included in McMurGut 1.1 MCMG754 by calculating percentage. The sum of the abundance reads from the covered genera were compared to the abundance reads from covered and non-covered genera, calculating again percentage.

### McMurGut 1.1 - ssniff diet creation

A skeleton medium was created by translating the ssniff chow composition chart (https://www.ssniff.com/) into a theoretical flux of metabolites available to the simulated microbiome of mmol per hour. Only those metabolites were considered that were part of the Gapseq metabolite list. It was determined that 5 g of chow per day is eaten, and 5.8 ml water is drunk a day. Mouse gut components such as water, mucin, urea and murine bile acids were included. Utilizing Micom’s complete_db_medium (Micom 0.35.0 in Qiime2-amplicon-2024.2) function, components were automatically added to the medium in aiming to enable growth of genera from MCMG754. Target doubling time was 0.1 (every ∼7h) with a maximum flux input of 10 mmol/hour. In the final composition, 132 of 134 genera were growing (check_db_medium). Oxygen and toxic substances, which were automatically added in a first completion step, were excluded in a second run (oxygen, formaldehyde, acetaldehyde, hydrogen peroxide, H2S, methanol, nitrite and hydrogen cyanide). Jupyter-scripts for this pipeline were made accessible at github (create_medium_v11.ipynb) [https://github.com/CME-lab-research/McMurGut].

### Metabolic simulation

Simulations were prepared in JupyterHub scripts and run in python (v3.8.15) via MICOM (v0.35.0) and Qiime2 (v2024.2). McMurGut model catalog MCMG754 on genus level, as well as taxonomy and abundance tables were loaded into Micom, running the build function with cutoff value at 0.0001, creating sample models and a manifest. Based on this, an optimal tradeoff value of 1.0 was determined with the tradeoff function. Microbial fluxes were created with the grow function, tradeoff 1.0, sample models, manifest and McMurGut murine diet file.

#### Differential mean flux

Microbial fluxes were processed with the functions prod_rates and compare_groups to find fold-changes and statistics (Mann Whitney test) differentially produced in cachexia.

#### Differential sum of flux per genus

To identify differential flux*abundance production in cachexia based on individual genera, microbial fluxes were processed via search_flux_species.ipynb. Briefly, metabolite flux and abundance from grow()-output set as export were multiplied to get flux*abundance, and stored for each metabolite, sample and taxon. All flux*abundances from samples from the same condition and taxon were added, leading to the sum of flux*abundance from a specific taxon under a specific condition. To calculate differential flux*abundance of a specific taxon between two conditions, flux*abundance of one condition (cachexia) was divided by the second condition (control), subtracted by 1, and values above 0 were increased in cachexia and below 0 were reduced in cachexia. If sample size of the two conditions was unequal, the condition with larger sample size was normalized to the condition with lower sample size (i.e. in E3, 5 controls to 4 cachexia, and in E123, 21 controls to 18 cachexia).

